# Method for quantifying the metabolic boundary fluxes of cell cultures in large cohorts by high resolution hydrophilic liquid chromatography mass spectrometry

**DOI:** 10.1101/2022.04.25.489416

**Authors:** Ryan A. Groves, Maryam Mapar, Raied Aburashed, Luis F. Ponce, Stephanie L. Bishop, Thomas Rydzak, Marija Drikic, Dominique G. Bihan, Hallgrimur Benediktsson, Fiona Clement, Daniel B. Gregson, Ian A. Lewis

**Affiliations:** University of Calgary, Department of Biological Sciences, Calgary, T2N 1N4, Canada; University of Calgary, Cumming School of Medicine, Department of Pathology and Laboratory Medicine, Calgary, T2N 1N4, Canada; Alberta Precision Laboratories, Calgary, T2L 2K8, Canada; University of Calgary, Cumming School of Medicine, Department of Community Health Sciences Calgary, T2N 1N4, Canada; University of Calgary, Cumming School of Medicine, Department of Medicine, Calgary, T2N 1N4, Canada

## Abstract

Metabolomics is a mainstream approach for investigating the metabolic underpinnings of complex biological phenomena and is increasingly being applied to large scale studies involving hundreds or thousands of samples. Although metabolomics methods are robust in smaller scale studies, they can be challenging to apply in larger cohorts due to the inherent variability of liquid chromatography mass spectrometry (LC-MS). Much of this difficulty results from the time-dependent changes in the LC-MS system, which affects both the qualitative and quantitative performance of the instrument. Herein, we introduce an analytical strategy for addressing this problem in large-scale microbial studies. Our approach quantifies microbial boundary fluxes using two zwitterionic hydrophilic interaction liquid chromatography (ZIC-HILIC) columns that are plumbed to enable offline column equilibration. Using this strategy, we show that over 360 common metabolites can be resolved in 4.5 minutes per sample and that metabolites can be quantified with a median coefficient of variation of 0.127 across 1,100 technical replicates. We illustrate the utility of this strategy via an analysis of 960 strains of *Staphylococcus aureus* isolated from blood stream infections. These data capture the diversity of metabolic phenotypes observed in clinical isolates and provide an example of how large-scale investigations can leverage our novel analytical strategy.

## Introduction

Metabolomics has emerged as a mainstream strategy for investigating cell physiology and understanding the molecular underpinnings of diseases. Liquid chromatography mass spectrometry (LC-MS) has become one of the most widely used platforms for metabolomics due to its relatively simple sample preparation requirements and its broad metabolite coverage^1^. Over the last 20 years, the depth of LC-MS methods has been expanded to capture ever broader transects of the metabolic network. These efforts have dramatically increased the scope of routine metabolomics^2^ and are now allowing researchers to map the previously unexplored “dark metabolome”^3^. Although these methods are indispensable for capturing the broadest possible scope of metabolism, not all metabolomics studies require this depth of coverage. This is particularly relevant in the context of large cohort studies, where the objective is to quantify a select set of molecules consistently across a large number of samples.

The analysis of metabolic boundary fluxes is one application where it is particularly important to have LC-MS methods optimized for long term stability, rather than depth of metabolite coverage. Boundary fluxes refer to the rates at which metabolites are consumed or secreted by cells^4^ and can be quantified by measuring changes in the composition of cell culture growth media over time. Quantifying boundary fluxes can provide a comprehensive understanding of cellular function by enabling flux balance models of cell metabolism to be optimized^5^, nutritional dependencies to be mapped^6^, and metabolic networks of model organisms to be optimized for industrial applications^7,8^. Moreover, we have recently shown that metabolic boundary fluxes can be used as a clinical microbiology assay for identifying microbial species and measuring antibiotic susceptibility profiles^9^. One critical distinction between boundary flux analysis and routine metabolomics is the complexity of the samples. Whereas intracellular metabolic phenotypes affect thousands of molecules^10^, cell media can contain 100 or fewer organic molecules^11^. The long chromatographic gradients that have been developed for intracellular metabolomics are not optimized for the modest complement of molecules found in culture media and needlessly increases the analysis time, cost, and error arising from time-dependent changes in LC-MS response factors.

Another key distinction between boundary flux studies and routine metabolomics is scale. Although relatively few metabolites need to be quantified in cell media, the long-term quantitative stability of these analyses is critical for minimizing batch effects across large cohorts. A variety of chromatographic^12^, ionization^13^, and other instrument factors contribute to the loss of LC-MS performance over time. One major factor is the deposition of lipids, proteins, salts, and other insoluble materials on the electrospray ionization source and ion optics^14^. These deposits cause progressive changes in the response factors that generally result in a loss of ionization efficacy. A related problem is the progressive deterioration of the chromatography column, which results from degradation of the stationary phase^12^. The time-dependent loss of chromatographic and ionization performance on LC-MS instruments is exacerbated in metabolomics studies because the complex samples can quickly foul the instrument. In contrast, the cell culture medium analyzed for boundary flux studies tends to be less complex and less chemically diverse than cell extracts, thereby enabling the analysis of larger cohorts of samples than would be practical for many metabolomics studies. However, the hydrophilic nature of most media components means that they are best resolved by hydrophilic interaction liquid chromatography (HILIC)^4,15,16^, which requires long equilibration times between injections to stabilize the stationary phase^17,18^. These long delays between injections are counterproductive in large cohort studies, when the primary objective is to maximize sample throughput. Novel methods that mitigate these sorts of shortcomings are thus necessary for large-scale analyses of metabolic boundary fluxes.

In summary, analyzing boundary fluxes provides a powerful approach for investigating cell physiology in large cohort studies, but modern metabolomics methods are not optimized for this application. Whereases metabolomics methods are tuned to maximize breadth of chemical diversity and depth of metabolite coverage, boundary flux studies track small collection of aqueous metabolites, have less complex input samples, and emphasize quantitative stability across large cohorts. To address this discrepancy, we developed a zwitterionic HILIC (ZIC-HILIC) LC-MS method that is specifically optimized to enable quantification of microbial boundary fluxes. Our approach uses sample injection multiplexing to minimize column re-equilibration times and a standardized LC-conditioning workflow for stabilizing the performance of the instrument. We show that this approach enables over 397 metabolites to be resolved in a 4.5-min gradient. Moreover, we show that metabolites observed in *Staphylococcus aureus* cultures can be quantified over 1,100 injections with a median coefficient of variation of 0.127 across the observed metabolites.

## Experimental Section

### Analytical standard preparation

For all qualitative work, the mass spectrometry metabolite library service (MSMLS™; Sigma-Aldrich) was used to prepare standards for determining chromatographic performance. Standards were received pre-weighed in 96-well plates with 5 µg in each well and were subsequently reconstituted to a concentration of 25 µg/mL using 50% methanol (1:1 Optima™ LC-MS grade water and methanol; Thermo Fisher Scientific). Following this, individual standard solutions were then combined equally into 65 pooled mixtures with either 12 compounds, or 6 compounds with 6 equivalents of 50% methanol so that all mixtures reached a final concentration per metabolite of 2.08 µg/mL. These mixtures were assigned such that they excluded isobaric compounds to minimize any possible cross interference when determining retention characteristics. Retention times were then annotated manually based on the peak signal intensity maxima in extracted ion chromatograms (+/− 10 ppm) for all observed [M+H]^+^ and [M−H]− ions with a signal intensity greater than 1e^4^. Pooled standard mixtures were run in both positive and negative ion mode and all manufacturer provided data (chemical name, elemental composition, mass characteristics, associated database entries), sub pool mixture number assignments, and observed retention characteristics were recorded in Table S1. Summary information of the number of compounds detected in each ionization mode was also tabulated in Table S2 based on this data.

For all quantitative work, a mixture of 85 commonly observed compounds was prepared in 50% methanol at varying concentrations (Table S3) and stored at –80 °C in 1 mL aliquots. Samples were then thawed on the day of use and a 16-point standard curve was prepared by serial dilution (see Table S3 for concentrations). Each metabolite concentration was calibrated to ensure that the metabolites quantified were within the linear range of response factors for each target metabolite. These samples were used in the determination of lower limit of detection (LLOD) and lower limit of quantification (LLOQ) values in positive and negative ion mode, which was defined as the lowest observed concentration in this standard curve with a signal ratio >3x and >10x, respectively, greater than the signal observed in a 50% methanol blank injection. Compounds observed in this data that were not present in the MSMLS™ set are annotated in Table S1. Some experiments in this work used additional 8- or 6-point standard curves where appropriate and are annotated in Table S3.

### Biological samples

Clinical isolates of *S. aureus* from patients with blood stream infections were collected from Calgary (Alberta, Canada) hospitals over a 27-month period by Alberta Precision Laboratories. 960 *S. aureus* isolates were identified following Clinical and Laboratory Standards Institute (CLSI) guidelines, and single colonies isolated from microbial cultures were stored as glycerol stocks. All activities were approved by the Conjoint Health Research Ethics Board (CHREB) of the University of Calgary under certificate REB17-1525.

In order to achieve uniform growth of selected *S. aureus* cultures, microbes were inoculated in 96-well plates filled with Mueller-Hinton broth originating from a single batch and were cultured overnight. Samples for all stages of growth were cultured aerobically at 37 °C in a humidified incubator with a controlled 5% CO_2_, 21% O_2_ atmosphere, without shaking. These were then sub-cultured and grown to an OD between 0.35–0.4, and culture supernatants were then fixed in methanol (1:1 v/v). Samples were centrifuged for 5 min at 4,000 x *g* and cleared supernatants were retrieved and stored at –80 °C. On the date of LC-MS analysis, stored supernatants were further diluted 1:10 with 50% methanol. Genomic sequencing, as described previously^19^, was performed on these cultures to verify microbial species and to screen for cross contamination.

### Liquid chromatography parameters

Chromatographic separation was accomplished on two Syncronis™ ZIC-HILIC columns (inner diameter, 2.1 mm; length, 100 mm; particle size, 1.7 µm) using a UHPLC Vanquish™ Integrated biocompatible system (Thermo Fisher Scientific) equipped with an auxiliary vanquish UHPLC pump to perform the analytical gradient while the main pump performs sample deposition and offline column re-equilibration. Mobile phases consisted of solvent A (20 mM ammonium formate, pH 3.0 in Optima™ LC-MS grade water; Thermo Fisher Scientific) and solvent B (Optima™ LC-MS grade acetonitrile with 0.1 % formic acid; Thermo Fisher Scientific). Solvent A was constituted by addition of ammonium formate salt followed by pH adjustment through titration with formic acid until a stable pH of 3.0 (+/− 0.05) was reached overnight. For each experiment, all required solvents were produced in one large batch to minimize the possibility of solvent batch effects. All gradients were run at a flow rate of 0.6 mL/min with a sample injection volume of 2 µL using this binary solvent system. Column temperature was maintained in a sealed compartment at 30 °C (+/− 0.005 °C).

During LC-MS analysis, two chromatographic gradients were employed simultaneously using the auxiliary vanquish UHPLC pump. This setup also utilized two sets of multiport switch valves (MSV) to direct alternating sample plugs to each column and manage sample analyte elution into the mass spectrometer (Figure S1). For the auxiliary pump, the following gradient was used with respect to solvent B: 95 %, from 0–0.5 min; 95 % to 5 %, from 0.5–3.5 min; 5 %, from 3.5–4.5 min. Concurrently, the main pump gradient used was: 95 %, from 0–0.5 mins; 95 % to 5 % from 0.5–0.75 min; 5 % from 0.75–2 min; 5 % to 95 % from 2–2.25 min; 100 % from 2.25–4.5 min. The lower MSV controlled sample introduction to each column and switched at the 0.4 min mark to allow sample plug deposition on the column head before switching to the auxiliary pump to begin the analytical gradient. The upper MSV switched between columns at the beginning of each run to direct the flow from the column currently eluting an analytical gradient into the mass spectrometer, and the equilibrating column to the waste.

### Mass spectrometry parameters

Electrospray ionization was undertaken at a sheath gas flow rate of 35, auxiliary gas flow rate of 15, sweep flow gas rate of 2 (all rates in arbitrary units), and all gas flows provided were high purity N_2_ gas. A spray voltage of −2.5 kV was used in negative ion mode and +3.0 kV was used in positive ion mode. Capillary temperature was maintained at 275 °C, and the auxiliary gas heater was set at 300 °C. A Thermo Fisher™ Q Exactive™ HF Hybrid Quadrupole-Orbitrap™ mass spectrometer was used to acquire data in negative and positive ion full scan mode, within a range of 50–750 *m/z*, at a resolution power of 120,000. The automatic gain control target was 1e^6^, and the maximum injection time was 200 ms.

### LC-MS system conditioning

To assess the long-term effects of chromatographic drift and column conditioning, the performance of a previously unused pair of chromatographic columns was monitored under standardized conditions over the course of 3,192 injections. To track the performance across this process, two types of samples were repeatedly injected: a 6-point standard curve (Table S3) and a pooled *S. aureus* sample made from 160 culture supernatants mixed at an equal ratio. These samples were collected on a repeating schedule wherein the 6 analytical standards were collected followed by 100 injections of the *S. aureus* pooled sample. Each sample was injected onto both columns, with injections continuously alternating between them until each column had reached a total of 1,500 *S. aureus* samples (3,192 total injections across the two columns, including standards).

### Boundary flux analysis of S. aureus clinical isolates

Our methods for quantifying boundary fluxes have been described in detail elsewhere^9^. Briefly, metabolites present in microbial cultures were collected prior to, and after incubating samples for 4 hours in Mueller-Hinton growth medium. Microbial culture media were diluted 1:20 (v:v final dilution) in 50% HPLC-grade methanol (Optima™ LC-MS grade methanol Optima™ LC-MS grade water; Thermo Fisher Scientific) and were analyzed by LC-MS. An 8-point analytical standard curve was collected interspersed between every 80 isolates and metabolite concentrations were calculated based on the observed response factors for each molecule. The amount of each metabolite produced/consumed over this period was then calculated.

### Peak Integration and statistical analysis

Extracted ion chromatograms for figures were generated in Thermo Scientific Xcalibur 4.0.27.19 software using a mass window of (+/−) 10 ppm. Peak assignment and quantification were performed using the open-source MAVEN software package^20,21^. Statistical analyses were performed using Python (Version 3) with seaborn and matplotlib packages and with an in-house R software package (https://zenodo.org/record/6403220#.Ykuh0mTF2_a). Retention factors (*k*) were calculated as shown in equation 1, where *t*_R_ is the observed retention time and *t*_*0*_ is the time required to clear the system void volume (0.42 min).

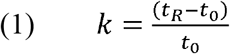

Coefficient of variation (CV) values were calculated as shown in equation 2, where σ is the standard deviation of LC-MS signals for a specific metabolite across the dataset and µ is the mean LC-MS signal for the same metabolite across the dataset.

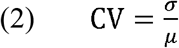

Metabolite signals were calculated as z-scores as shown in equation 3, where the signal for each metabolite (x) is shown relative to the mean signal for the same metabolite observed in Mueller Hinton media samples (µ) divided by the standard deviation (σ) of the same metabolite observed across the Mueller Hinton media samples.

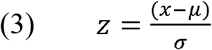

Graphically displaying z-scores of metabolite consumption/production is challenging because the data span several orders of magnitude and because z-scores include both positive and negative data. To display these data (Figure 5), enrichment factors (*EF*) were calculated for each metabolite signal. Enrichment factors were defined as shown in equation 4, where c was set to 10. This transformation is adapted from the Rocke & Durbin variance stabilization algorithm^22^.

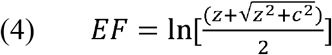

### Safety Considerations

Reagents used in this investigation do not pose any significant safety risks outside of those experienced regularly when working with moderate strength acids and solvents.

## Results and discussion

### Multiplexed chromatographic approach

Metabolomics methods are not optimized for analyzing low-complexity media samples to generate quantitatively stable readouts over thousands of injections. To address this, we developed a sulfobetaine-based ZIC-HILIC chromatography method for assessing microbial boundary fluxes that separates the media components of interest that are commonly found in microbial cultures. The stationary phase we selected was previously identified as one of the better performing chemistries for capturing broad coverage of aqueous metabolites^23,24^. Using this stationary phase, we evaluated 40 combinations of solvent elution profiles, pHs, organic solvents (acetonitrile, methanol), and buffer salts (ammonium formate, ammonium acetate, ammonium bicarbonate) to find a method that optimizes run time while maintaining broad metabolite coverage. These refinements were made using a mixture of 85 metabolite standards that are commonly found in cell culture media (Table S3). The optimized conditions we present here (Figure S1) use a 4.5-min linear chromatographic gradient with acetonitrile, using 20 mM ammonium formate at pH 3 (see *Liquid chromatography parameters* for details). Figure 1 shows 23 representative extracted ion chromatograms from our pooled standard set showing that the metabolites resolved using this method have an average peak width of ∼2.4 sec (Table S5) and are evenly distributed across the elution profile.

**Figure 1.**
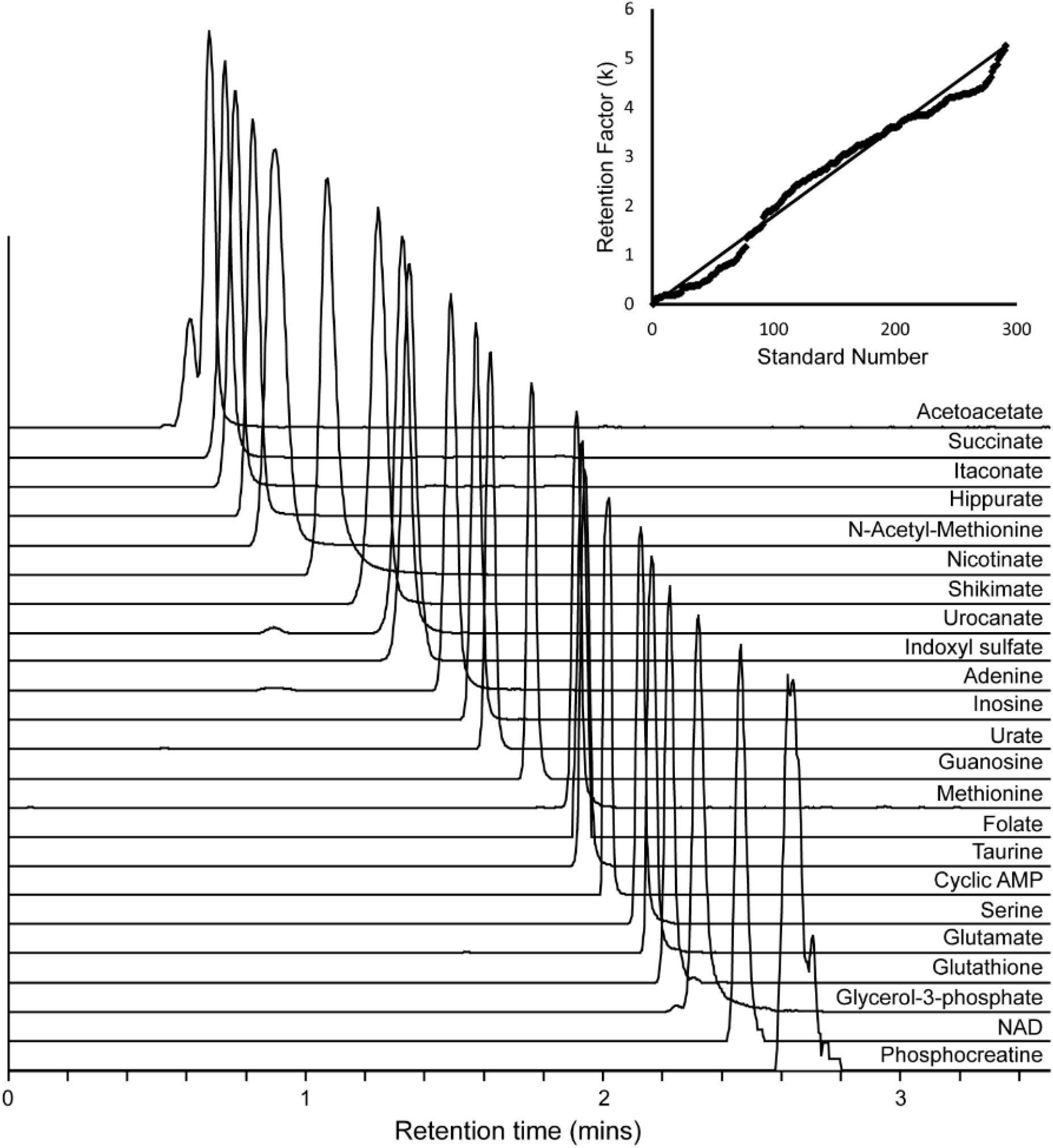
Chromatographic performance of a broad transect of metabolites resolved by our multiplexed ZIC-HILIC method observed in negative ion mode. Main panel displays stacked extracted ion chromatograms of 23 representative standards selected from the MSMLS™ library covering a diversity of metabolite chemistries of biological relevance, with optimal peak shapes. Observed signal intensities are normalized across the set for visual comparison without the interference of differential ionization effects. Inset displays the distribution of retention factors (*k*) of all 313 compounds detected in negative ion mode from the MSMLS™ library. The solid line indicates a theoretical perfect spacing of compounds across the run between the void volume and last observed standard retention time.

To assess the qualitative performance of this method, we analyzed a commercially available MSMLS™ library of metabolite reference standards containing 639 compounds (Table S1). These standards have representatives from a broad transect of polar metabolites including amino acids, nucleic acids, organic acids, energy metabolites, vitamins, and compounds important to cellular redox control. From this library, 397 compounds were ionizable and chromatographically resolvable. The chromatographic separation of these diverse analytes was close to the theoretical optimum peak spacing (Figure 1 inset).

One of the primary disadvantages of HILIC is that it frequently requires long re-equilibration times between injections^25^. To address this, we plumbed two HILIC columns into an LC-MS system using a secondary HPLC pump (Figure S1). This arrangement allowed columns to be reconditioned offline, and thereby compresses the overall analytical run times without sacrificing time on column re-equilibration. Using this strategy, we were able to reduce the total sample-to-sample cycle time to 4.5 min.

Another factor affecting the length of analytical analyses is that some metabolites do not ionize in both positive and negative mode. This problem can be addressed by collecting data in consecutive injections using the two modes but has the obvious disadvantage of doubling the analytical run time. This problem can be addressed on some LC-MS platforms using polarity switching methods^26^ wherein the source polarity is alternated between scans. However, the scanning rate on the instrument use in this study (Thermo Fisher Orbitrap, QE-HF) was not fast enough to adequately sample the narrow elution peaks we observed when using these polarity switching methods. However, our analysis of the MSMLS™ library indicated that 313 of the 391 detectable compounds (80%) could be analyzed in negative mode (Table S2). Given that the goal of our method was to maximize sample throughput rather than increase the breadth of metabolite coverage, we elected to only collect negative-mode data.

### LC-MS conditioning for large cohorts

To assess the quantitative stability of our method in large cohorts, we injected 1,500 pooled *S. aureus* samples alternately (3,000 injections in total) on a two previously unused columns. A 6-point pooled analytical standard curve was collected every 100 samples on each column (192 injections in total). Variation in the signals from 55 metabolites present in the *S. aureus* samples and 77 metabolites present in the standards mixture were then tracked across all 3,192 injections (Figure 2a) as well as retention time stability (Table S7). The overall performance of our method, as judged by run-to-run stability and progressively smaller technical errors, stabilized after ∼400 injections with incrementally smaller improvements observed thereafter (Figure 2a). After 378 injections, the median CV for all metabolites reached a threshold CV of 0.15 (median CV across injections 0-378 = 0.171; median CV across injections 379-1500 = 0.127). Given this behavior, we recommend 400 replicate injections of biological samples to fully stabilize the LC-MS system prior to the analysis of large cohorts using this method.

**Figure 2.**
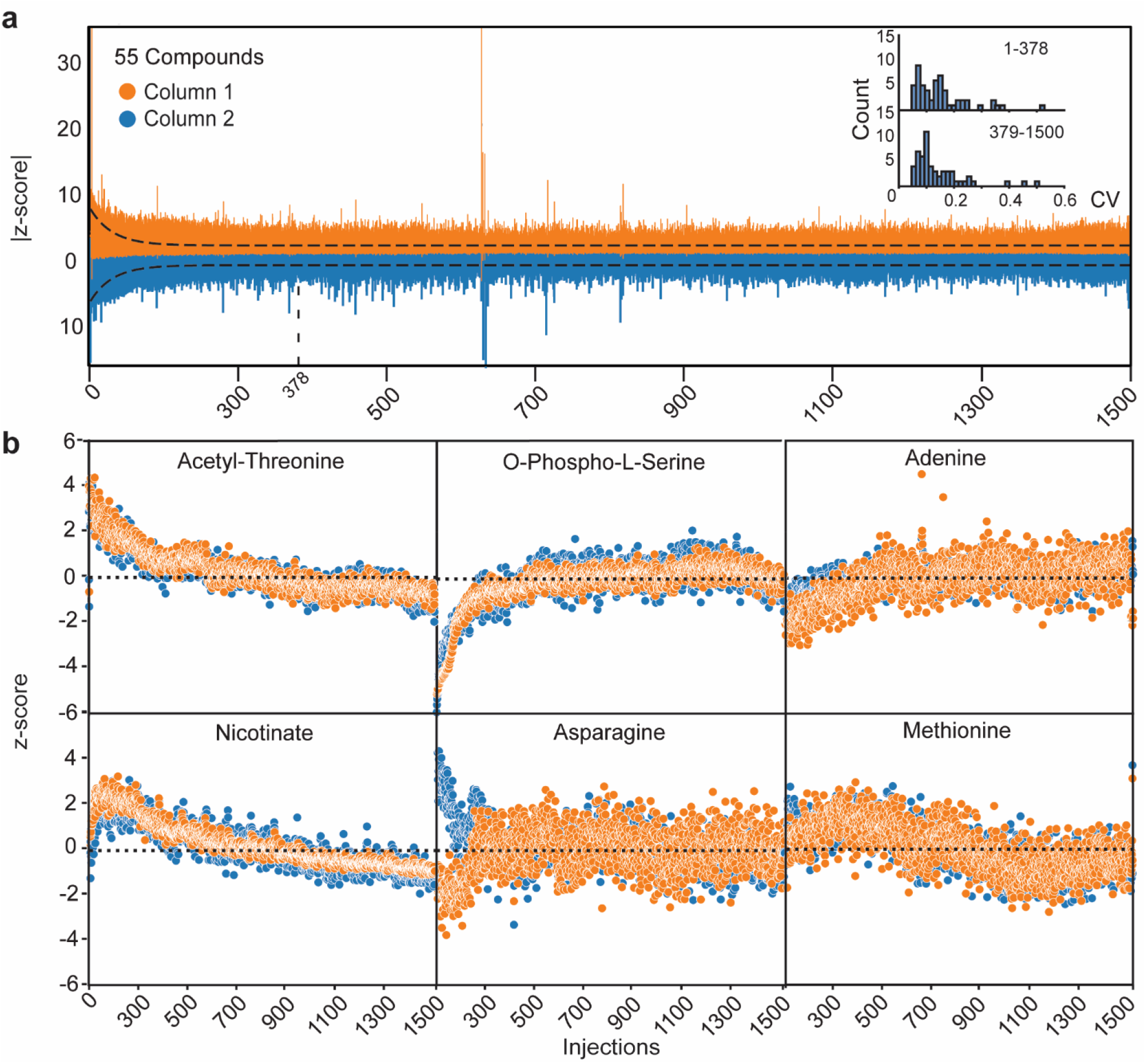
Quantitative stability of metabolites from *S. aureus* cultures over 1,500 injections. Variations in signal intensities were monitored for 55 metabolites using our multiplexed ZIC-HILIC MS analytical strategy. For this study, two new Syncronis™ ZIC-HILIC columns were used. Performance was monitored over repeated injections with a pooled *S. aureus* extracellular extracts (n = 3,000). Blue and orange points differentiate data originating from each of the two columns. (a) To illustrate the most variable signals for each injection, the z-score for each metabolite was computed (relative to the median signal and standard deviation observed across all injections) and the upper quartile of z-scores observed across all metabolites in each injection was computed. These data were then plotted for each column independently as absolute values and overlayed. The median z-scores observed for across metabolites were then fitted to a decaying exponential (dashed line). The point where CV values crossed the 0.15 point is noted (378). Inset shows a histogram of CV values observed across all detected metabolites from 1–378 sample injections and from 379–1500 sample injections. (b) Compound-to-compound difference and column-to-column differences were observed in the instrument response factors over the conditioning period. A representative selection of these differences is illustrated (shown as z-scores for each individual metabolite). The patterns for all 55 metabolites are provided in Figure S2.

Importantly, the conditioning pattern observed for each compound was not uniform: significant compound-to-compound differences were observed as well as some column-to-column to differences (Figure 2b, Figure S2). For example, phosphoserine and other phosphate-containing compounds showed a rapid increase in response factors over the first 400 injections, whereas acetyl-threonine and several other acetylated amino acids showed a decrease in response factors over the same period. Interestingly, several amino acids (asparagine, serine, glutamine, histidine, and lysine) showed inverse patterns between the two columns: compounds eluting from one column progressively gained signal intensity whereas the same compounds eluting from the other column progressively lost signal intensity. However, after the recommended 400 sample conditioning period, we did not observe any significant compound-to-compound changes in response factors: all signals followed a progressive loss of signal that is consistent with the slow accumulation of insoluble materials (salts, lipids, proteins, carbohydrates, etc.) on the column, source, and ion optics. Collectively, these data show that 400 injections are sufficient to stabilize CVs, minimize compound-to-compound differences in the behaviour of response factors over time, and standardize the behaviour of metabolites across the two chromatographic columns. After this conditioning period, we showed that response factors are stable over more than 1,100 injections (median CV = 0.127; Table S6).

### Sensitivity of multiplexed ZIC-HILIC MS

To assess the quantitative performance of *multiplexed ZIC-HILIC MS*, we prepared a 16-point standard curve (n = 6) using a mixture of 85 compounds. We then determined the LLOD and LLOQ for each compound (Figure 3; Figure S3; Table S4). The median LLOD across these compounds was 24.2 nM in negative mode and 61.0 nM in positive mode ionization. Median LLOQs were 39.1 nM and 98.6 nM in negative mode and positive mode, respectively. Given that the primary goal of this method is to enable quantification of abundant media compounds, which are generally present at micromolar to low millimolar concentrations, these sensitivities are more than adequate for quantifying microbial boundary fluxes.

**Figure 3.**
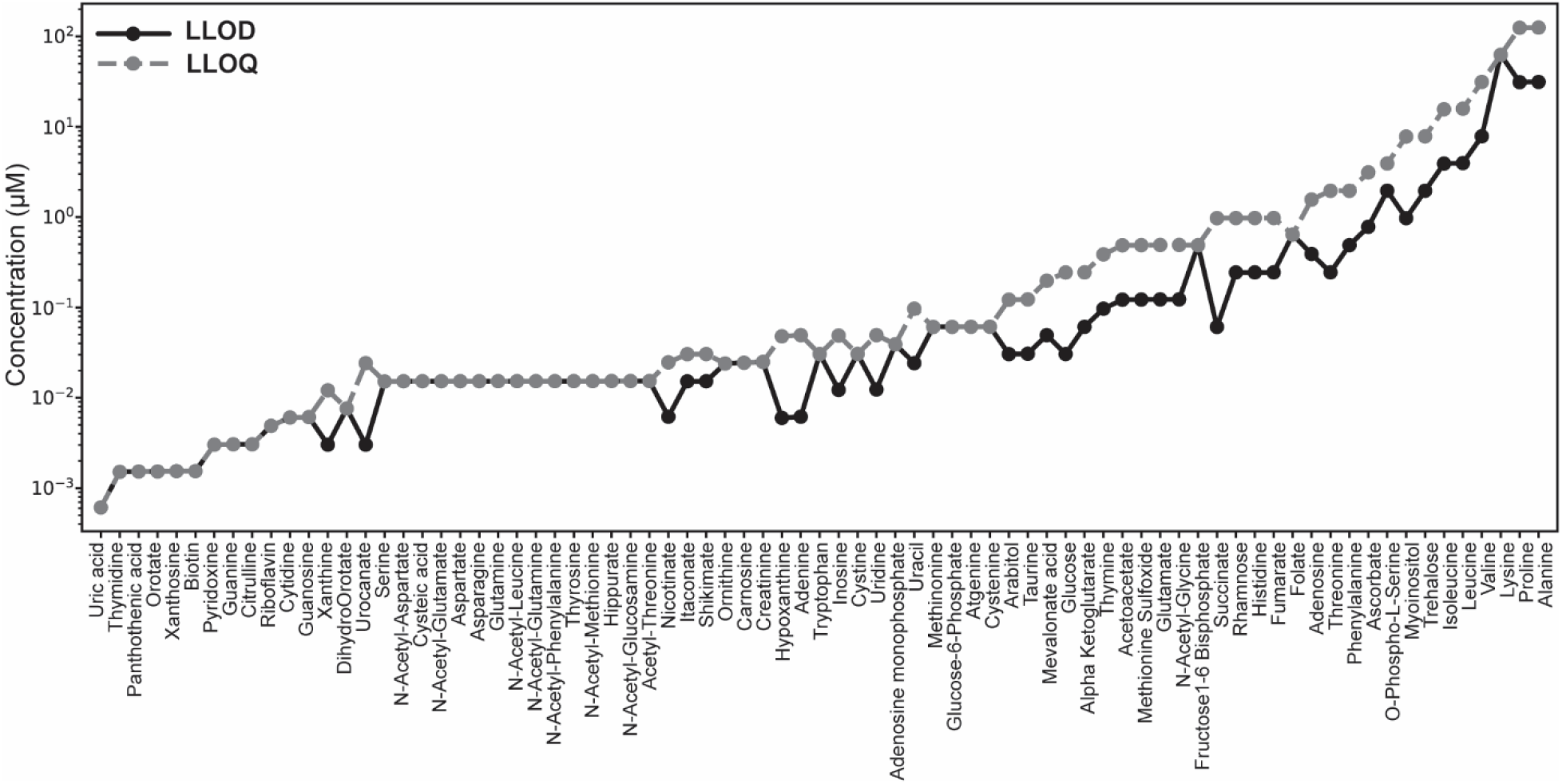
Sensitivity limits for our multiplexed ZIC-HILIC MS method. Median LLOD and LLOQ levels for 77 metabolites were computed from a 16-point pooled standard curve (n = 6) in negative ion mode. LLODs and LLOQs were defined conservatively as the lowest concentration metabolite standard whose signal was empirically observed to be >3 times, and >10 times the noise threshold, respectively. Similar data were observed in positive mode ionization (Figure S3), all LLOD/LLOQs are provided in Table S4.

### Analyzing metabolic boundary fluxes by multiplexed ZIC-HILIC

*S. aureus* is a common cause of blood stream infections^27,28^ and changes in its metabolism have been linked to virulence (e.g. via the arginine catabolic mobile element)^29^. Thus, metabolic profiling of *S. aureus* could potentially be harnessed as a tool for clinical diagnostics^9^. Currently, most *S. aureus* metabolomics studies have been conducted using a laboratory strains^30^ and relatively small collections of clinical isolates^31^. Currently, it is unclear if the metabolic phenotypes described in these studies represent the full spectrum of metabolic diversity found in populations of *S. aureus*. To better understand the metabolic diversity of *S. aureus* populations, and to illustrate the utility of multiplexed ZIC-HILIC MS, we analyzed the metabolic boundary fluxes of 960 clinical isolates of *S. aureus* collected from blood stream infections (27 months of infections in the Calgary area). Clinical isolates were grown in 12 batches of 80 samples and each batch was analyzed alongside an 8-point calibration curve of standards. Overall, we observed consistent metabolic phenotypes across the cohort (Figure 3), as judged by signal intensities from the mixtures of standards (median CV = 0.125, Table S8).

Our boundary flux analysis showed that *S. aureus* cultures preferentially consume glucose and nucleic acid-related compounds (uridine, xanthine, adenine, and adenosine) and secrete a broad range of metabolic products (mevalonate, citrulline, succinate, N-acetyl-methionine, thymine, orotate, and urocanate; Figure 5). These phenotypes were surprisingly homogenous across the cohort with only a few molecules (e.g. citrulline) showing evidence for strain-to-strain variability. Collectively, this phenotype is consistent with a metabolic model wherein *S. aureus* relies on glycolysis for carbon and energy needs and uses nucleotide salvage pathways when the appropriate precursors are present in the environment. The highly stable nature of the phenotypes we observed is somewhat surprising given the prevalence of auxotrophy that has been reported previously in the literature^33,34^. However, our use of a rich microbial growth medium may provide sufficient nutritional diversity, allowing each strain to satisfy their basal biomass and energetic needs with minimal disruption in the overall flow of molecules into and out of the cells.

**Figure 4.**
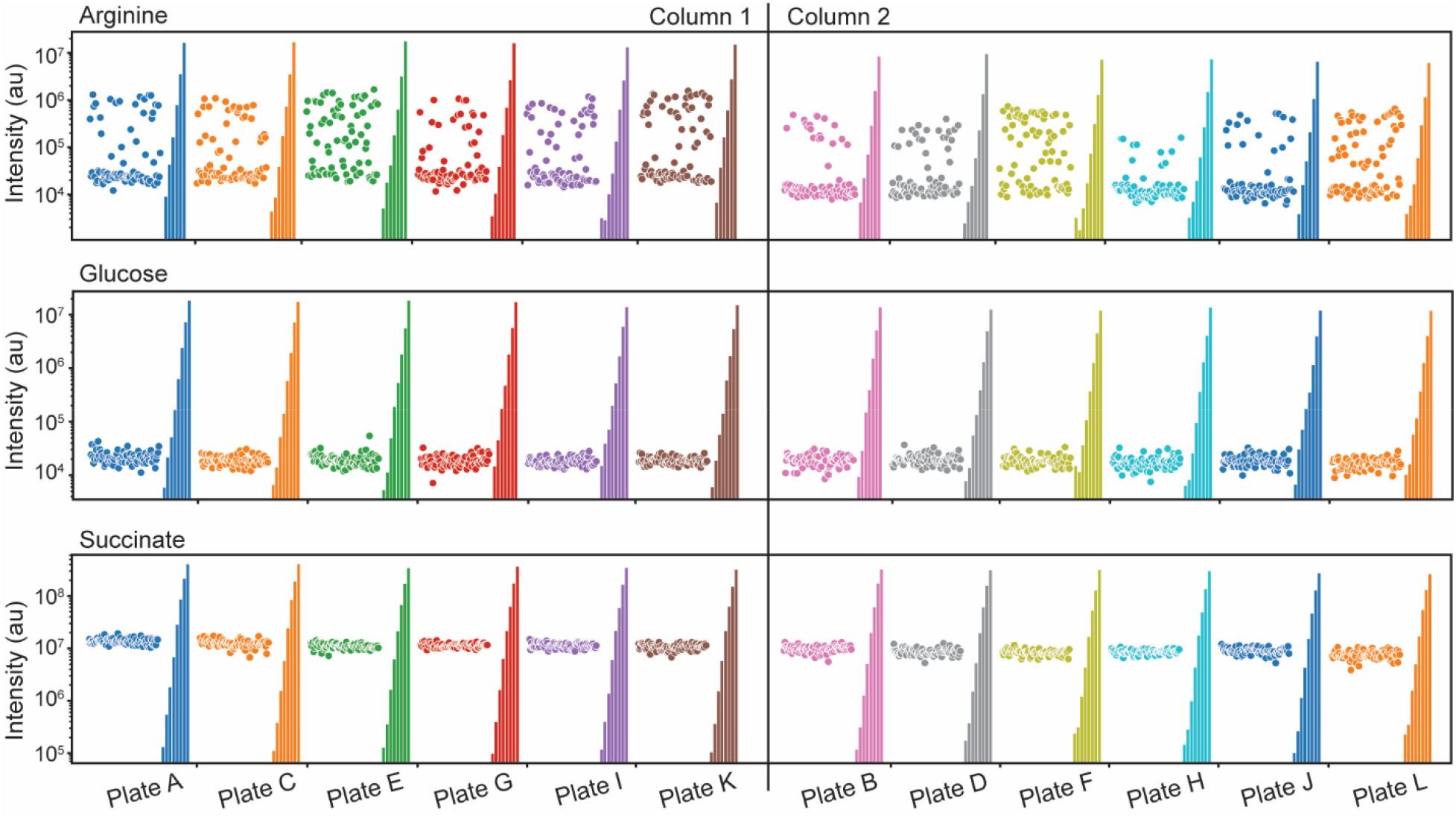
Metabolic phenotypes observed via multiplexed ZIC-HILIC. (a) Metabolite signals observed across a cohort of 960 *S. aureus* culture extracts originating from clinical isolates. Arginine, glucose, and succinate were chosen as biologically significant metabolites that are relevant to microbial phenotypes. The variability in arginine shown here is clinically relevant given that the arginine catabolic mobile element (ACME) is linked to virulence^29,32^.

**Figure 5.**
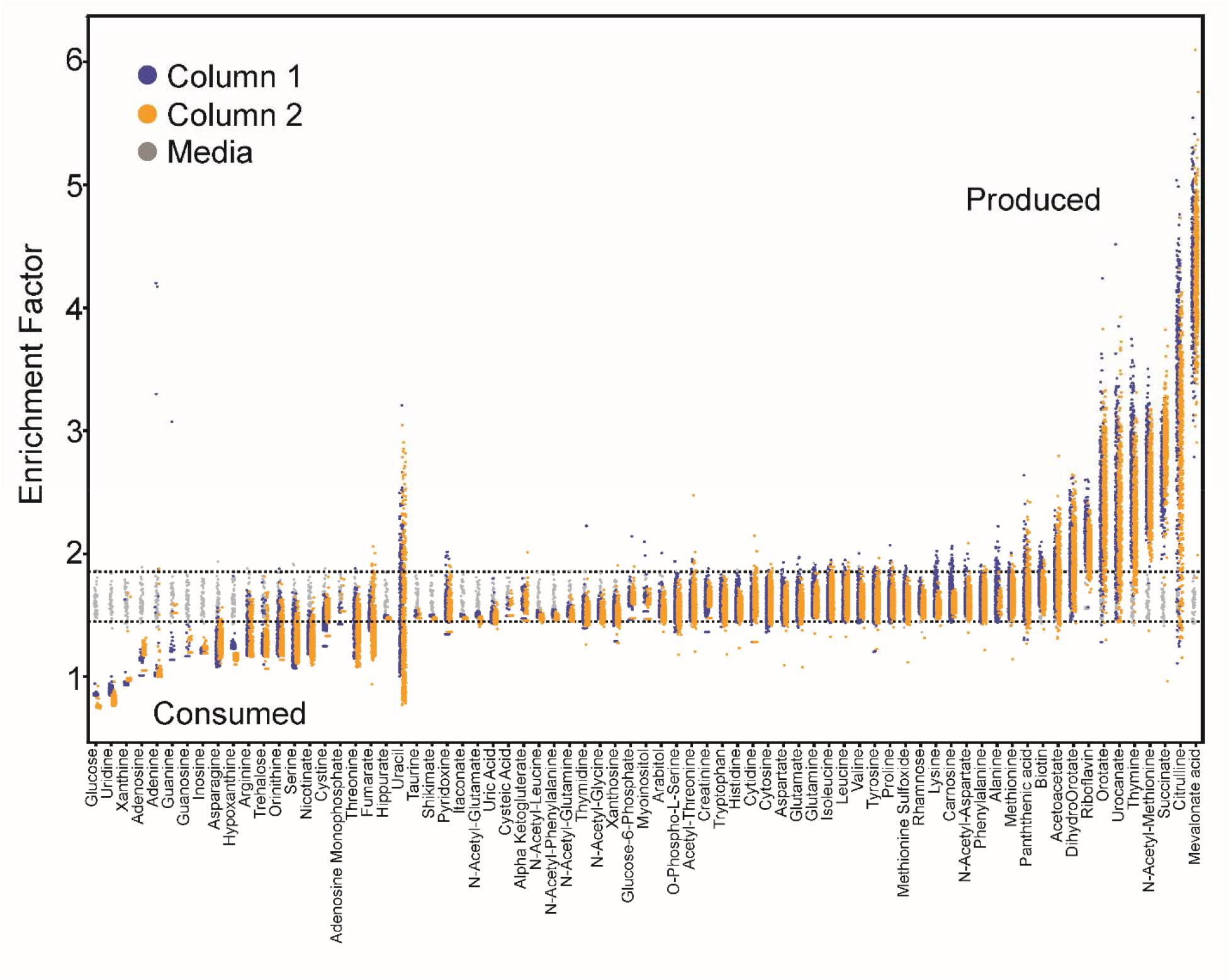
Metabolomic boundary flux of *S. aureus* clinical isolates. LC-MS analysis of 77 metabolites levels observed in *in vitro* cultures of 960 *S. aureus* isolates from a 27-month collection period of bloodstream infections in Calgary. Isolates were grown under standardized conditions in Mueller-Hinton broth. Orange/blue points indicate metabolite values observed in individual isolates that were run on each of the two analytical columns. Dashed lines denote the two standard deviation boundaries of metabolite signals observed in the media control. Enrichment factors are a log-transformed z-score (unitless) calculated according to equation 4 in the experimental section. This transformation enables the data, which span many orders of magnitude in z-score space, to be visualized. Raw data are provided in Table S9 prior to transformation.

Interestingly, three of the metabolites that do show evidence for metabolic diversity across the clinical cohort are arginine and two of its catabolites, ornithine and citrulline. Changes in the boundary fluxes of these compounds are consistent with the presence/absence of the arginine catabolic mobile element (ACME)^32^, a virulence-associated trait that is selected in tandem with the methicillin resistance encoding SCC*mec* operon in the USA300 clone^29^ that has been reported in the Calgary population^35^. Detecting these phenotypes in clinical cohorts highlights one potential application of the analytical strategy introduced here.

The method we introduce here can be applied to a range of applications where the primary goal is to understand the rates at which a discrete set of microbial metabolites are excreted to or consumed from the culture medium. These applications include 1) constructing/refining flux balance models^4,5^, 2) screening large libraries of isolates for auxotrophy or other metabolic variability^33,34^, 3) as a tool to accelerate bioengineering projects, particularly those that are seeking to use saturation mutagenesis and other high-throughput methods to maximize metabolic outputs in microbial systems^7,8^, and 4) as a strategy for rapidly screening inhibitor libraries to identify new antimicrobial lead compounds. Moreover, we have recently shown that microbial boundary fluxes can be used as a diagnostic readout for differentiating microbial pathogens and measuring their antibiotic susceptibility profiles^9^. In summary, the method we present here has broad applicability to any large cohort study of *in vitro* culture systems where the primary objective is to quantify the rates at which metabolites are consumed or secreted to the culture medium.

### Limitations in the multiplexed ZIC-HILIC MS strategy

Although the methods introduced here enable quantification of boundary fluxes in large cohorts, they have some inherent limitations that should be considered in the context of each study. One important consideration is that this method relies on a custom hardware configuration involving two HPLC pumps and two switch ports, a setup which is not standard on most LC-MS instruments. In addition, we show that conditioning the LC-MS using 400 injections of biological samples is needed to stabilize the performance of the method. This conditioning period would be disruptive if conducted on the main LC-MS platform; however, conditioning could be completed offline on a separate HPLC. Beyond these hardware considerations, the most significant limitation of the method is its relatively limited sensitivity. Modern metabolomics methods are pushing ever further into secondary metabolism with higher sensitivity methods.

The method introduced here is not suited to these in-depth applications and will not cover the chemical diversity, the low abundance molecules, and the isomer resolution that is possible with longer methods. One specific shortcoming we have identified is with our multiplexed ZIC-HILIC MS strategy is that pyruvate, lactate, and short chain fatty acids are not detected – all of which are important for comprehensive mapping of metabolic fluxes. Despite these limitations, we have shown that the method can be applied to large microbial cohorts and introduces minimal quantitative variability across thousands of samples.

## Conclusions

Metabolomics has emerged as a mainstream strategy for understanding the molecular underpinnings of life. Modern LC-MS metabolomics methods can capture over a thousand molecules and a broad transect of central carbon metabolism. Despite this, there are relatively few methods that have been optimized for large cohort, boundary flux studies involving thousands of samples. The multiplexed ZIC-HILIC MS methods introduced here addresses this gap and enables consistent quantification of microbial metabolic boundary fluxes. We show how this strategy can be used to map metabolites flowing into and out of cells and stably quantify these phenotypes across thousands of clinical samples. This study provides a template for other large-cohort metabolomics studies and provides a comprehensive set of reference data for implementing this method in other laboratories.

## Associated Content

### Supporting Information

Supplemental Information.pdf – PDF document containing: Figure S1 “Schematic overview of the HPLC plumbing configuration for a multiplexed two column chromatography system”, Figure S2. “z-score based chromatographic drift analysis from repeated pooled *S. aureus* sample injections”, and Figure S3. “Quantitative positive ion mode LLOD/LLOQ performance”.

TableS1.xlsx – Excel document containing Table S1. MSMLS qualitative compound database.

TableS2.xlsx – Excel document containing Table S2. MSMLS summary data.

TableS3.xlsx – Excel document containing Table S3. Analytical standard curve concentrations.

TableS4.xlsx – Excel document containing Table S4. Overall quantitative method performance in negative and positive ion mode.

TableS5.xlsx – Excel document containing Table S5. Chromatographic peak widths of 23 selected chemical compounds.

TableS6.xlsx – Excel document containing Table S6. Coefficient of variation values from *S. aureus* column conditioning study.

TableS7.xlsx – Excel document containing Table S7. Retention time shift analysis during *S. aureus* conditioning study.

TableS8.xlsx – Excel document containing Table S8. CV performance of standards accompanying 960 *S. aureus* sample cohort.

TableS9.xlsx – Excel document containing Table S9. Raw signal intensity data for 960 *S. aureus* isolates.

## Author Information

### Author Contributions

The manuscript was written through contributions of all authors. All authors have given approval to the final version of the manuscript.

## Acknowledgments

This work was supported by funding from Genome Canada, the Canadian Institute of Health Research, the Government of Alberta, Alberta Precision Laboratories, and Alberta Innovates through Genome Alberta under its 2017 Large Scale Applied Research Project and 2016 Genomic Applications Partnership Program competitions; the Canadian Institute of Health Research (162790); the University of Calgary (Biomedical Engineering 10011121); and Alberta Precision Laboratories. I.A.L is supported by an Alberta Innovates Translational Health Chair. T.R. is supported by an Eyes-High Postdoctoral Fellowship from the University of Calgary (10011121). Metabolomics data were acquired at the Calgary Metabolomics Research Facility, which is supported by the International Microbiome Centre and the Canada Foundation for Innovation (CFI-JELF 34986). This work was made possible in part by a research collaboration agreement with Thermo Fisher.

---

For Table of Contents Only

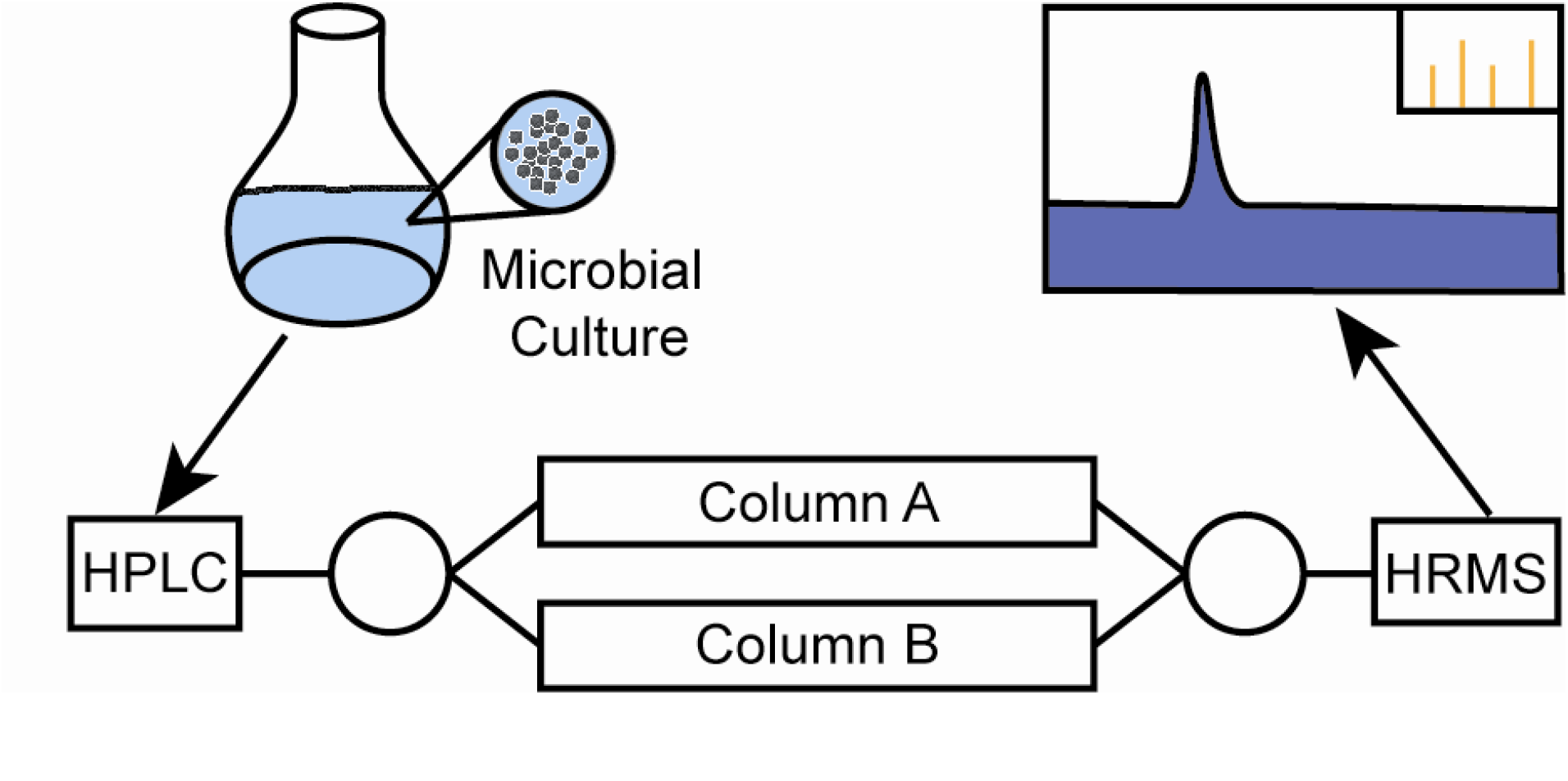

