## Supplemental Information for "Method for quantifying the metabolic boundary fluxes of cell cultures in large cohorts by high resolution hydrophilic liquid chromatography mass spectrometry"

**Table of contents**

Figure S1……………………………………………………………………………………….........S2

Figure S2………………………………………………………………………………………..S3

Figure S3………………………………………………………………………………………..S4

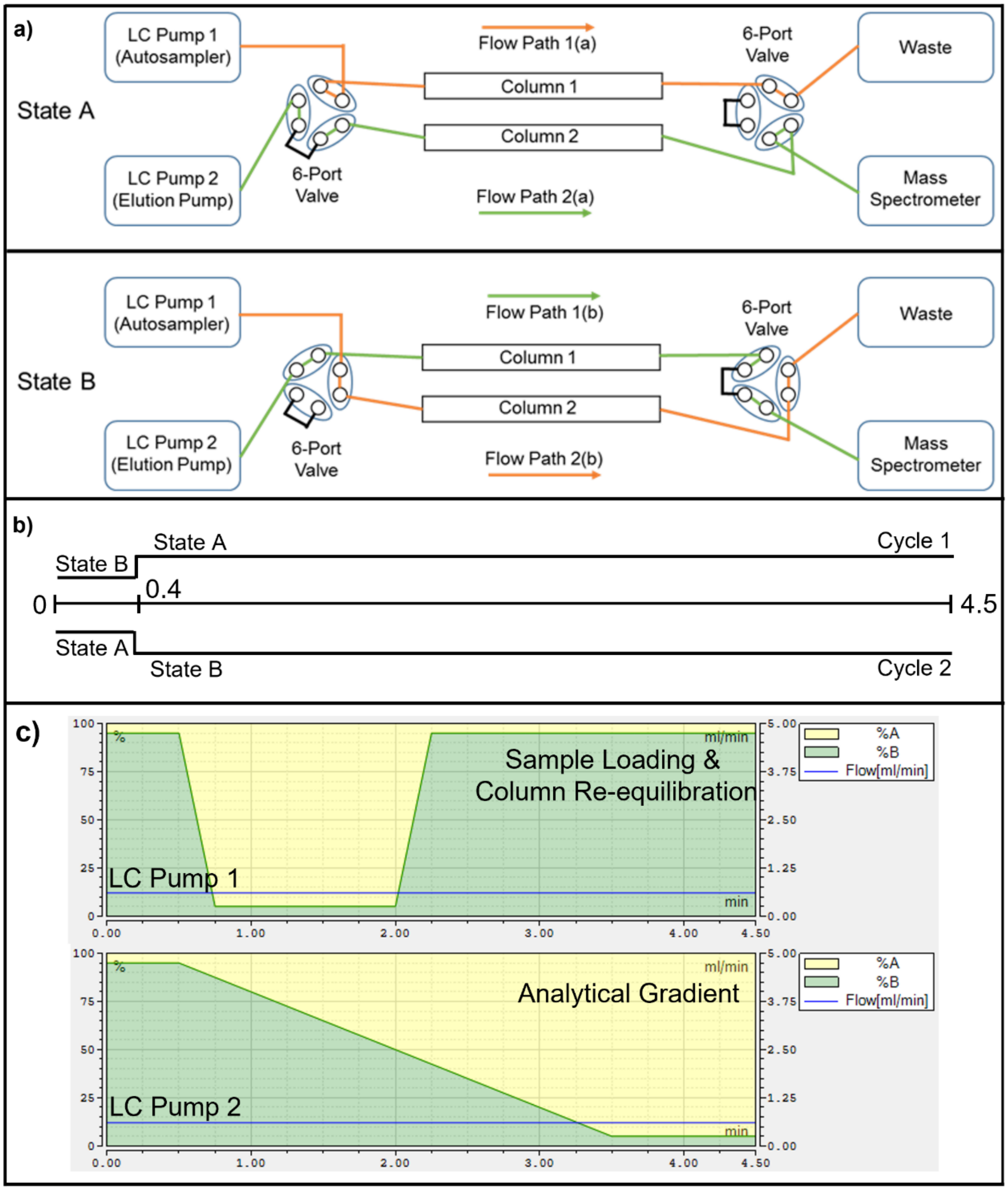


**Figure S1. Schematic overview of the HPLC plumbing configuration for a multiplexed two column chromatography system.** (a) Plumbing schematic of the two states used so that the system toggles between the two analytical cycles employed for this method. (b) Timing of the two states used when running samples; samples are run by alternating between cycle 1 and cycle 2 continuously. (c) Chromatographic gradients run by LC pump 1 and LC pump 2. Yellow/green colouring represents the percentage of solvent A and B respectively. LC pump 1 handles sample injection followed by flushing and re-equilibration of the offline column after switching out of line with the mass spectrometer at the 1-minute mark. LC Pump 2 performs the analytical gradient on the inline column following sample deposition on the head of the column under strong binding conditions (95% acetonitrile). Note that these gradients do not change between cycle 1 and cycle 2, only the columns which they are connected to via state A/B.

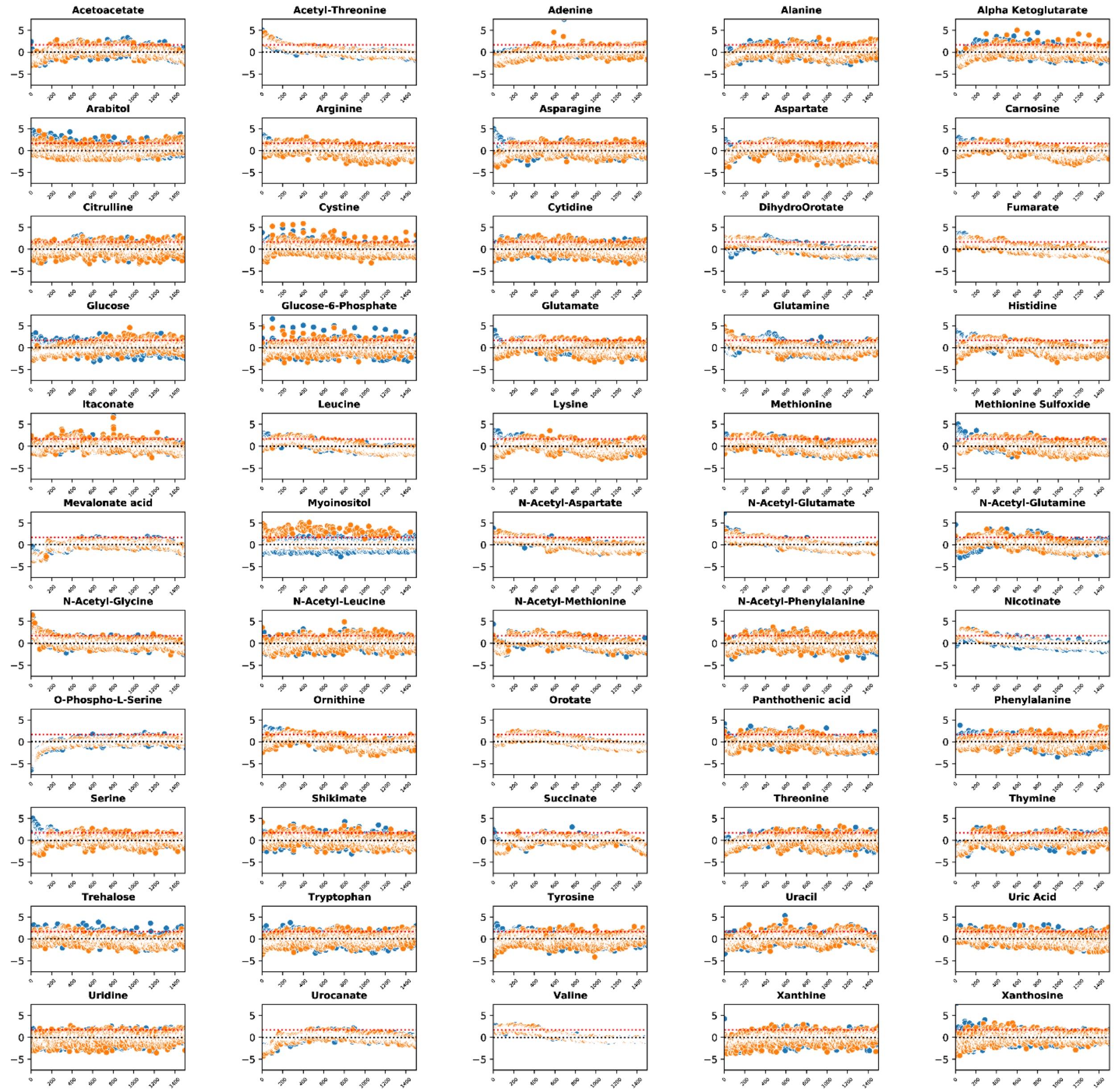


**Figure S2. z-score based chromatographic drift analysis from repeated pooled *S. aureus* sample injections.** Calculated z-scores for all detected compounds observed from 3000 injections of a pooled *S. aureus* sample across two columns displayed individually. Blue and orange colours denote samples injected on each individual chromatographic column.

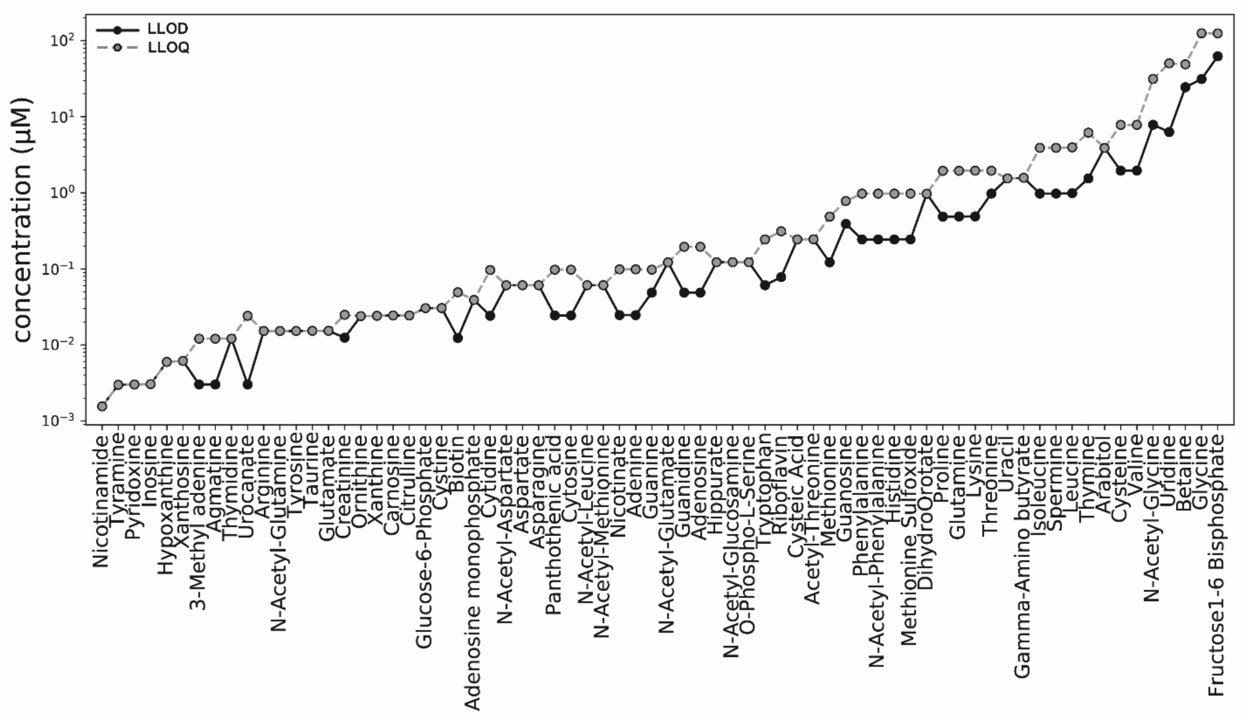


**Figure S3. Quantitative positive ion mode LLOD/LLOQ performance.** Median limit of detection and limit of quantitation levels for all observed metabolites from a 16-point pooled standard curve in positive ion mode (*n* = 6). Exact sample concentrations described in Table S3.
